# Show me your neighbours, and I’ll tell you what you are – cellular microenvironment matters

**DOI:** 10.1101/231282

**Authors:** Timea Toth, Tamas Balassa, Norbert Bara, Ferenc Kovacs, Andras Kriston, Csaba Molnar, Lajos Haracska, Farkas Sukosd, Peter Horvath

## Abstract

To answer major questions of cell biology, it is essential to understand cellular complexity. Modern automated microscopes produce vast amounts of images routinely, making manual analysis nearly impossible. Due to their efficiency, machine learning-based analysis software have become essential tools to perform single-cell-level phenotypic analysis of large imaging datasets. However, an important limitation of such methods is that they do not use the information gained from the cellular micro- and macroenvironment: the algorithmic decision is based solely on the local properties of the cell of interest. Here, we present how various microenvironmental features contribute to identifying a cell and how such additional information can improve single-cell-level phenotypic image analysis. The proposed methodology was tested for different sizes of Euclidean and nearest neighbour-based cellular environments both on tissue sections and cell cultures. Our experimental data verify that the microenvironment of a cell largely determines its entity. This effect was found to be especially strong for established tissues, while it was somewhat weaker in the case of cell cultures. Our analysis shows that combining local cellular features with the properties of the cell's microenvironment significantly improves the accuracy of machine learning-based phenotyping.

## Introduction

Recent improvements in microscopy and computational cell biology have led to an explosion of data volume, often as large as millions of images. These large bioimaging datasets raised a strong need for automated and objective analysis tools^1^. Various software (both commercial and open-source) have been developed^2–4^ for image and computational data analysis. One of the most commonly used open-source software is CellProfiler^5^. It has modules for various image processing tasks that can be performed sequentially to form a pipeline. Via this pipeline, biological objects, usually nuclei, cytoplasm, and cells can be identified, and metric features of these objects such as area, shape, texture, and intensity can be calculated. Recent studies propose segmentation solutions for the distinguishing of even more complex shape morphologies such as touching^6^ or overlapping^7^ cells.

Despite their advantages, single-cell segmentation approaches often prove to be inefficient, for example in the case of tissue section image analysis. Therefore, we have decided to use the simple linear iterative clustering (SLIC) superpixel segmentation method for the analysis of tissue sections as described in this article. Superpixel algorithms group pixels into larger coherent regions, therefore, they often replace the conventional pixel grid algorithms nowadays^8^. They have become increasingly popular in computer vision applications recently because they are fast, easy-to-use, and produce high-quality segmentations. The SLIC algorithm creates superpixels by clustering pixels according to similarities in intensity and proximity in the image plane^9^.

Machine learning methods are designed to learn functional relationships from examples based on features rather than from manual verification of entire experiments^10^. Compared to conventional approaches, these methods are more efficient in handling multi-dimensional data analysis tasks such as distinguishing phenotypes that are defined by a high number of features^11,12^. CellProfiler Analyst is an extension to CellProfiler and performs supervised learning from extracted features to recognize a single phenotype in individual cell images^13,14^. CellClassifier allows researchers to view the original microscope images so the observer can annotate an individual cell in its natural context^15^. Enhanced CellClassifier is another approach based on CellProfiler metadata, suitable for multi-class classification^16^. This program enables the differentiation between complex phenotypes. Advanced Cell Classifier (ACC) is a graphical image analysis software tool that offers a variety of machine learning methods^17^. CellProfiler Analyst 2.0 has been released recently and has many advantages compared to its previous version^18^. It is written in Python, works with multiple machine learning methods, can perform cell- and field-of-view-level classification, and has a visualization tool to overview an experiment. ACC 2.0 includes phenotype finder, a novel method to automatically discover new and biologically relevant cell phenotypes^19^. Additionally, some software are capable of classifying whole images instead of objects within images (e.g., WND-CHARM, CP-CHARM)^20,21^.

An important limitation of the above-mentioned software is that they work at the single-cell level only: they do not derive data from the micro-, or the macroenvironment of the cell; therefore, they do not take the population context of the cell of interest into account. It has been shown that single-cell heterogeneity in cell populations is determined by both intrinsic and extrinsic factors^22–24^. Based on previous studies on genetically identical single cells, we are convinced that the diversity in their phenotypic properties is defined by the features of growing cell populations that inherently create microenvironmental differences to which cells finally adapt^25,26^. Cells of tissues are also not organized randomly: the basis of the cellular landscape is formed as early as during the differentiation process, which is determined by well-established biological mechanisms. Therefore, the cellular milieu strongly determines single-cell entity. Thus, it seems reasonable to use the environmental data of each single cell of interest for machine learning applications.

In this paper, we present a systematic analysis of how cellular microenvironment affects the phenotypic analysis of single cells using supervised machine learning. Aggregated features of the environment were calculated for different neighbourhood sizes, and machine learning recognition rates were compared. Various popular machine learning methods were used for the evaluations. The methodology was tested on cell culture and tissue section data. Our results show that by incorporating the properties of the cellular microenvironment into phenotypic analysis tools, we can largely outperform classical approaches.

## Materials and methods

### Datasets

To test our hypothesis on real biological and clinical data, we chose two different datasets. First, we tested our hypothesis on data of a cell-based breast cancer cell line treated with different drugs used in clinical practice. Next, images of a thin section of urinary bladder cancer tissue were analysed to verify our results and examine the performance of the novel method tested.

#### MCF-7 High-Content-Screening Dataset

The first dataset we used is a publicly available MCF-7 (MCF-7) breast cancer cell line set (available online at the Broad Bioimage Benchmark\ Collection^27^ (https://www.broadinstitute.org/bbbc/BBBC021/) that had been treated for 24 hours with 113 various small molecules at eight different concentrations. Briefly, the treatments applied on the MCF-7 breast cancer cell line included a distinct set of targeted and cancer-relevant cytotoxic compounds that induced a broad range of gross and subtle cell phenotypes. Next, the cells were fixed, labelled for DNA, F-actin, and ?-tubulin and were imaged by fluorescent microscopy. Images included in the publicly available dataset were taken from 55 microtiter plates of 96-well format 55 plates of a 96-well microtiter plate. This dataset comprises about 39.000 images containing approximately 2 million cells^28^. For our analysis, we used the singlecell phenotypic annotation presented by Piccinini and colleagues^19^. Nine phenotypic and a debris class were identified, and approximately 1500 cells were labelled (see Supplementary Table S1).

#### Urinary bladder cancer tissue sections

Our second image dataset comprises images of a urinary bladder cancer (UBC) tissue. Hematoxylin-eosin (HE) staining of slides of the urinary cancer tissue was carried out according to the routine histopathologic process. Briefly, formalin-fixed and paraffin-embedded tissues were cut in 4-μm-thick sections and stained in a Tissue-Tek DRS 2000E-D2 Slide Stainer (Sakura, Nagano Japan) instrument according to the manufacturer’s instructions. Images were taken by an Axio Imager Z.1 (Carl Zeiss, Jena Germany) microscope equipped with an EC Plan-NEOFLUOAR 20x/0.5NA lens using the AxioVision SE64Rel.4.9.1.1 (Carl Zeiss, Jena Germany) software. This dataset contains 38 images. We distinguished eight phenotypic classes and labelled an average of 1200 superpixels for each superpixel size (see Supplementary Table S1).

### Evaluation software

For the experiments, we used an image analysis and machine learning software (SCT Analyzer) developed by Single-Cell Technologies Ltd. (Szeged, Hungary). This software is a newly developed visual tool designed to be user friendly; it offers massive parallelization and provides an out-of-the-box solution for a wide-range of cell-based and histological analyses. It supports all the best-known operating systems (Windows, Mac, Linux). It can handle standard image formats such as JPEG, TIFF, and PNG and supports microscopy and high-content screening formats both at the image and at the metadata levels. Versatile image pre-processing (illumination correction, filtering) and cell segmentation methods (including SLIC segmentation) are implemented. An interactive interface helps the user to annotate segmented regions into an arbitrary number of phenotype classes. Numerous machine learning methods are available even for single-cell-level prediction. Last but not least, an active learning interface is provided to maximize user efficiency^29^.

#### Segmentation

Images from the high-content-screening dataset of drug-treated MCF-7 samples were segmented with a custom version of CellProfiler 2.2.0 (Fig. 1a). Nuclei were detected using the adaptive Otsu algorithm. Nuclei touching the boundaries and cells smaller than 5 μm were discarded. Cytoplasm of cells was extracted using adaptive thresholding with watershed separation based on the nuclei as seed points.

**Figure 1.**
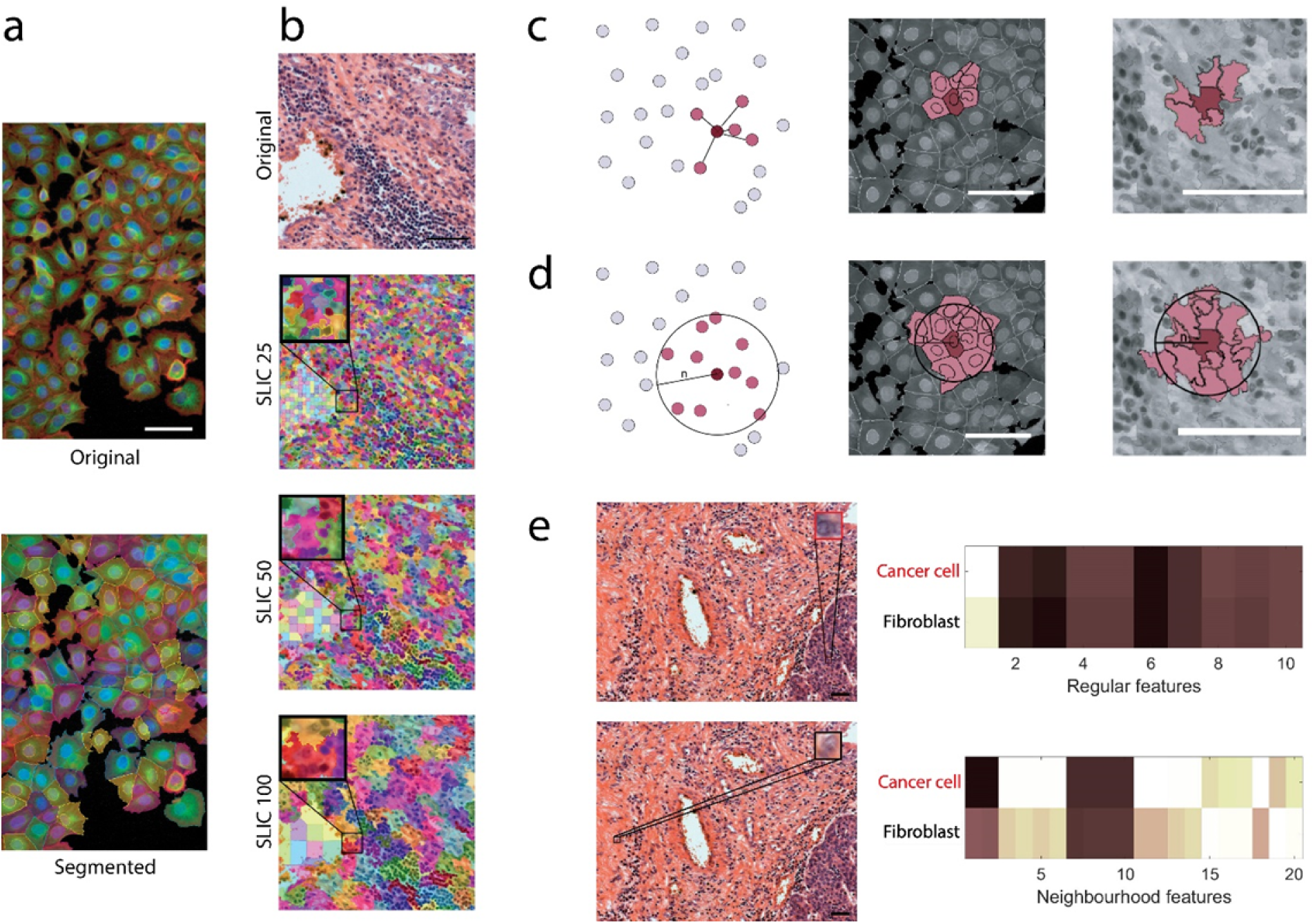
Segmentation and feature extraction. **(a)** Segmentation of the MCF-7 breast cancer cell line using CellProfiler 2.2.0. Scale bar: 50 μm **(b)** SLIC superpixel segmentation of urinary bladder cancer tissue section images. Images were segmented into superpixels of different sizes (25, 50, 100 pixels). Scale bar: 50 μm **(c)** The K-nearest neighbours (KNN) method, illustrated in a schematic figure and in real cell culture and tissue section scenarios, K=5. Scale bars: 25 μm **(d)** The n-distance method, illustrated in a schematic figure and in real cell culture and tissue section scenarios, n=50 pixels (cell culture: 19.51 μm, tissue sections: 13.5 μm). Scale bars: 25 μm **(e)** Superpixels containing two different phenotypes (cancer cell and fibroblast) share highly similar regular features, but features of their neighbourhoods differ significantly. Scale bars: 50 μm.

In the case of the UBC section images, the SLIC superpixel segmentation algorithm was used^9^. We set different superpixel sizes: 25, 35, 50, 75, and 100 pixels - 6.75, 9.45, 13.5, 20.25, and 27 μm (Fig. 1b). In all cases, we forced connectivity between superpixels if a superpixel was smaller than 20, 25, 40, 60 or 75 pixels, respectively.

#### Feature extraction

The most commonly used cell-based and neighbourhood features were extracted. Regular features describe the intensity, texture, and shape of individual objects. A full list of these features can be found in Supplementary Note 1. In this study, we analysed single-cell or superpixel image analysis results using only regular and neighbourhood features and the combination of these.

To represent the properties of the microenvironment, we assumed that cellular features had already been calculated for all cells/superpixels. The center of mass was measured for each segmented area and was used as a reference point for distance calculation. We used two different approaches to get the neighbours of a cell or superpixel: the K-nearest neighbours (KNN) and the N-distance methods (Fig. 1c, d). For the KNN method (where ‘K’ stands for a positive integer), we selected the K-nearest neighbours for each cell/superpixel based on Euclidean distance. For the distance-based approach, we took a fixed (n pixel) distance-based radius around an area’s reference point and selected all cells/superpixels within this range. At the end of this selection process, we had all neighbouring cells/superpixels for each individual superpixel. From that point, we used the same methodology for both the KNN and the N-distance methods to calculate neighbourhood features.

Neighbourhood features were derived from the mean, median, standard deviation, minimum, and maximum statistics of cellular features. Distance statistics (mean, median, standard deviation, minimum, maximum) describing the localization of neighbours were computed. For the Euclidean distance-based approach, an extra feature was calculated describing the extent of the neighbourhood, i.e., the number of neighbours within the given range (in the case of KNN, this number was known since it was the ‘K’ value).

Figure 1e represents an example in which two cells of different phenotypes share very similar basic features and local appearance, however, considering additional neighbourhood features, an obvious difference is revealed between them.

#### Machine learning

After feature extraction, we used the SCT Analyzer system to create the annotated single-cell set for machine learning classification. In the case of the high-content-screening dataset, we distinguished nine phenotypic classes: abundant, rounded, elongated, multinucleated, bundled microtubule, peripheral cytoskeleton, punctate actin foci, decreased cell size, and fragmented nucleus (Fig. 2a) and a debris class. The entire list of labelled cells in each class can be found in Supplementary Table S1. We paid special attention to avoid annotating identical cell types in close proximity and thus the biasing of neighbourhood features (because in this case cells would have extremely similar features and one may favourably bias the evaluation). In general, most of the images contained no more than one annotated cell. The performance of the annotated set was imported to Weka 3.8.1 (http://www.cs.waikato.ac.nz/ml/weka/), a machine learning and statistical framework.

**Figure 2.**
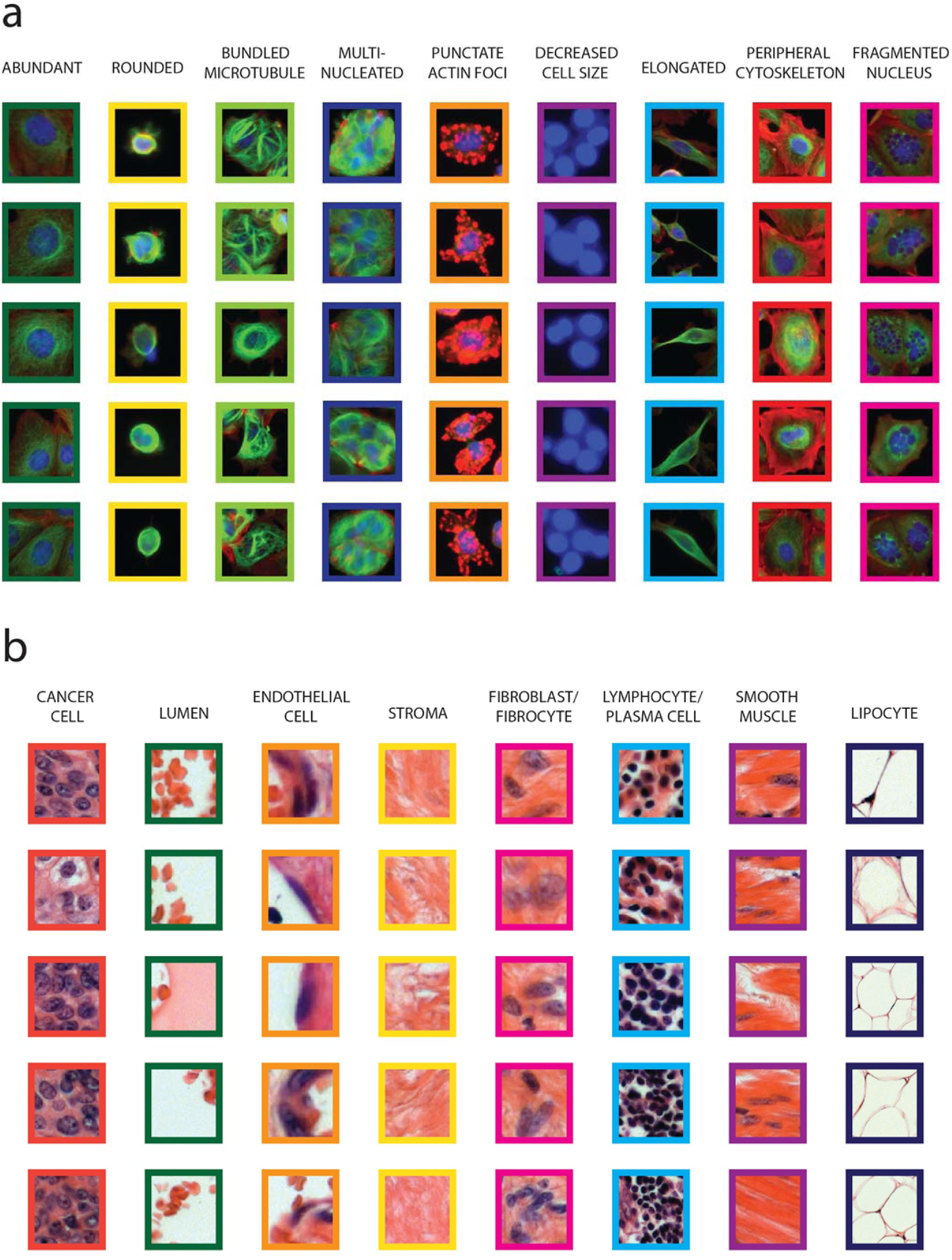
Distinguished phenotypes. **(a)** Cells of nine different phenotype classes identified in the MCF-7 High-Content-Screening Dataset. **(b)** Eight phenotypic classes in the UBC tissue image dataset.

Within the UBC cancer tissue image dataset, we distinguished eight different phenotypic classes: cancer cell, lumen, endothelial cell, stroma, fibroblast-fibrocyte, lymphocyte-plasma cell, smooth muscle, and lipocyte (Fig. 2b) and a debris class. The entire list of labelled superpixels in each class can be found in Supplementary Table S1. As this dataset included less images than the previous one, it was unavoidable to have annotated cells close to one another. However, to make sure that the cells in close proximity land either in the training or in the test set, so that we can prevent a potentially positive influence of evaluation, we performed the fold generation for cross-validation at the image level instead of the cell level. This way, folds of images were created and were used for the 10-fold cross-validation measurement.

For the high-content dataset, we investigated the size of the neighbourhood only, while in the case of the UBC image set, we also evaluated the classification performance as a function of superpixel size.

In both cases, we evaluated five different classification methods: the Weka’s Naïve Bayes, the Random Forest, the Support Vector Machine (SMO), the Logistic Regression (Simple Logistic), and the Multilayer Perceptron approaches. We used 10-fold cross-validation to measure the classifiers’ performance.

## Results

We evaluated the performance of neighbourhood features on both image sets. During this analysis, we compared different machine learning techniques as well as the extent of the neighbourhood and whether local features, microenvironmental features, or the combination of these features give the best results. Based on strong advice from biologists and pathologists, who definitely highlight the importance of cellular environment, we expected that taking neighbourhood features into account will increase the performance of machine learning. This speculation was further supported by our observation that cells in different phenotypic classes can share highly similar local properties, but the extended features unambiguously distinguish them (Fig. 1e). We also speculated that there should be an optimal (nonzero, but not extremely large) extent of the neighbourhood's size where classifiers perform best (i.e., an optimal neighbourhood size with respect to accuracy).

### Improved accuracy in cell culture

In the case of the MCF-7 breast cancer cells, we used 5-25 nearest neighbours for the KNN, and neighbours ranging between 100 and 1200 pixels (39.025-468.3 μm) for the Euclidean distance-based analysis. Cross-validation results show that increasing the number of neighbours improves accuracy (Fig. 3a) for all of the classifiers. Interestingly, we did not observe the expected tendency, i.e., accuracy did not peak, but increased and reached a plateau with a higher number of neighbours. The best performance we observed was for the 800-pixel distance (312.2 μm) radius using the SMO classifier. In this case, accuracy reached 88.57%, which is 8% better than that achieved when considering local features only.

**Figure 3.**
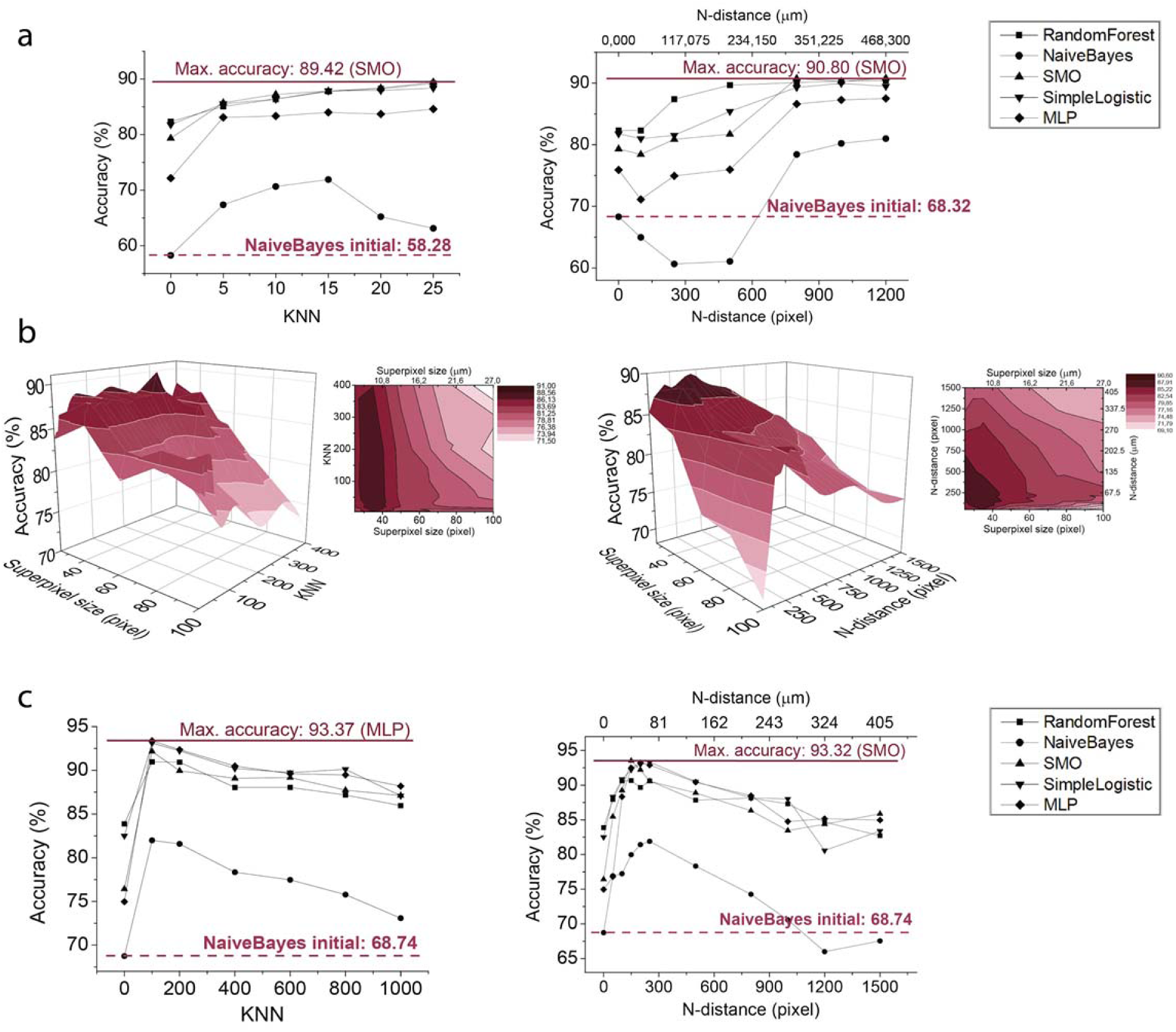
Comparison of the performance of machine learning methods (RandomForest, NaiveBayes, SMO, SimpleLogistic, MultilayerPerceptron) on different neighbourhood distances. **(a)** Machine learning accuracies in the cell culture dataset using neighbours selected with the KNN (left) and the N-distance methods (right). (We note that principal component analysis was performed before the Naïve Bayes and the Multilayer Perceptron calculations to reduce computational complexity.) **(b)** Three-dimensional (3D) illustration (and its contour) of the performance of the RandomForest algorithm in the UBC tissue dataset with respect to different superpixel and neighbourhood sizes using the KNN (left) and the N-distance (right) methods. **(c)** Machine learning accuracies on the best performing superpixel size (SLIC35, based on Figure 3b results) in the case of the UBC tissue image dataset.

### Neighbourhood features have major influence on phenotyping tissue sections

In the case of the UBC tissue images, we analysed neighbourhood features from 5-3000 nearest neighbours for the KNN method and between 100- and 1500-pixel distances (27-405 μm) for the Euclidean neighbourhood method. First, we tested the RandomForest algorithm for all training sets. Cross-validation results for different SLIC sizes are plotted in Figure 3b, indicating that we observed the expected tendency for machine learning accuracy. The best performance appears at superpixel size 35 (when a superpixel region is approximately 89.3 μm^2^) in the case of 100-nearest neighbours; at this superpixel size this means that we calculate with features of the microenvironment from an average of 137.15 μm radius. In this case, accuracy reaches 90.96%, while using only regular features at the same superpixel size and same KNN value, accuracy is only 83.87%, equalling to over 7% increase in performance. We also examined other supervised classification models at superpixel size 35 (Figure 3c). MLP classifier was found to produce the best accuracy (93.37%) when we used 100-nearest neighbours. Without the neighbourhood features, accuracy was only 74.96%. A similar tendency was detected for the other classifiers tested. In each case, higher accuracy was reached using neighbourhood features compared to considering regular features alone.

## Discussion

Phenotypic single-cell analysis has utmost importance for basic biological discoveries and next-generation digital pathology evaluations. The comparison of neighbouring cells, at lesser or larger distances on the slide, is an important part of the routine histological work. For example, the recognition of anisocytosis or anisonucleosis, two cellular hallmarks of malignancy, based on the subjective morphological comparison of neighbouring tumour cells is of great importance. Until now, digital pathology evaluation strategies focused on single cell features, and to the best of our knowledge, the environment of individual cells was beyond its focus.

Here, we present a machine learning-based phenotyping method that combines local and neighbourhood features, and we demonstrate that taking cellular and tissue component neighbourhood into account significantly increases recognition accuracy (Fig. 3). This improvement was detected both for cell cultures and tissue sections, although we should note that it proved to be somewhat weaker (but still prominent) in the case of cell culture images. This difference can be explained from a biological viewpoint: human tissue is a collection of similar cells, which acquire larger similarities during differentiation than that visible among cultured cells. Consequently, they predict each other’s morphology more than the loose colony of cells. Evidence is visible in Figure 1e, which displays two morphologically overlapping normal fibroblast and urothelial cancer cells. Regular features themselves cannot differentiate them; the proposed neighbourhood features can. Predictions and confusion matrices presented in Figure 4 confirm the effectiveness of neighbourhood features in machine learning calculations. Employing the information gained from the cellular microenvironment increases the ability of machine learning methods to identify patterns between cells near each other.

**Figure 4.**
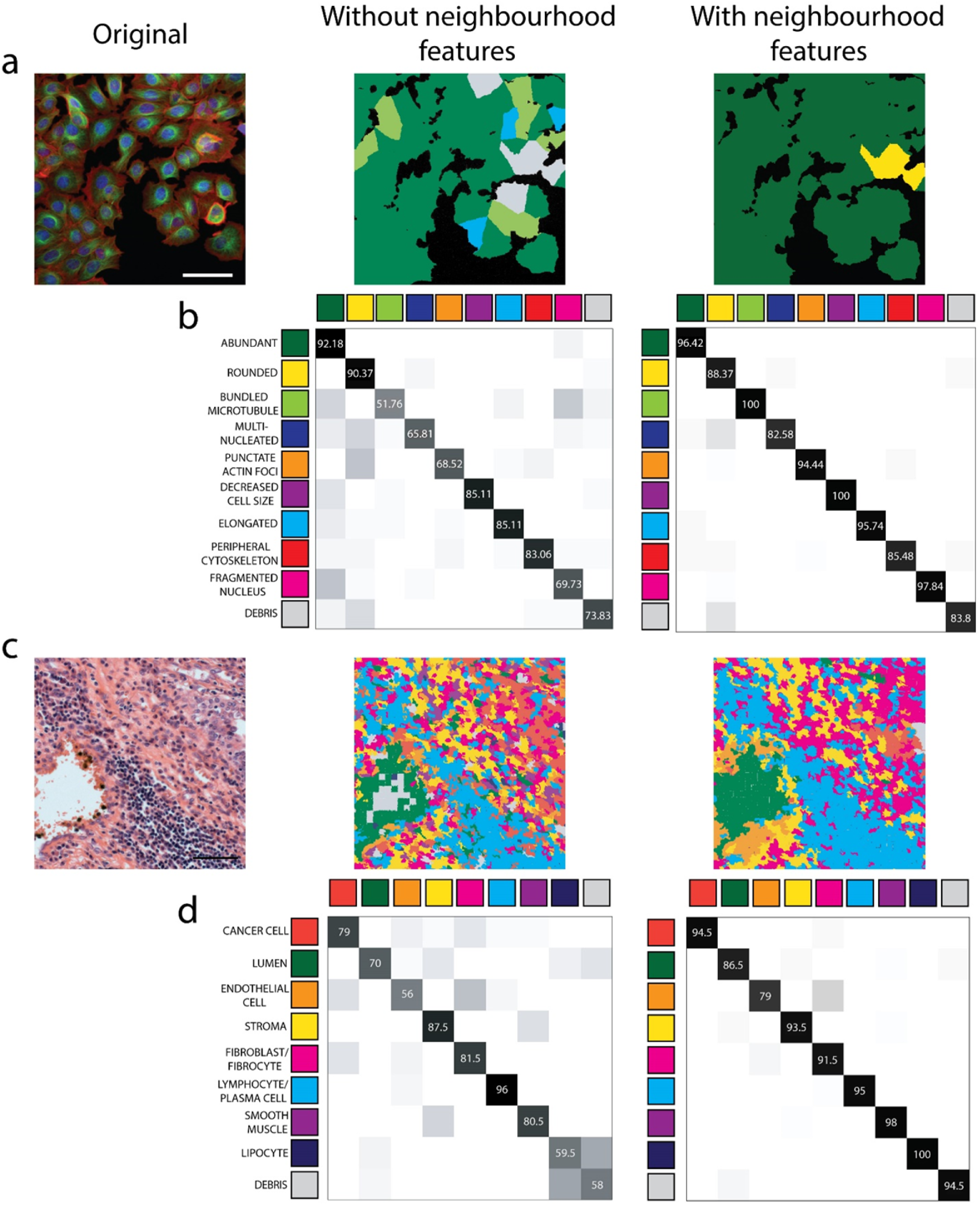
The effect of taking cellular microenvironment into account. **(a)** Prediction based on machine learning (SMO) in the cell culture dataset. Original image (left), scale: 50 μm, prediction using regular features only (middle), prediction using regular and neighbourhood features (right) **(b)** Confusion matrices of the best machine learning performance (SMO) in the MCF-7 breast cancer cell dataset, taking the features of single-cells into account (middle) and considering neighbourhood features (N-distance: 1200 pixels, 468.3 μm) as well (right) **(c)** Prediction based on machine learning (MLP) in the UBC tissue dataset. Original image (left), scale: 50 μm, prediction using regular features only (middle), prediction using the combination of regular and neighbourhood features (right), superpixel size: 35 pixels (9.45 μm) **(d)** Confusion matrices of the best machine learning performance (MLP) in the case of the tissue section images. Calculations using base features only (middle) and taking cellular microenvironment into account (KNN, K=100).

The most exciting questions are: how many neighbours have to be called and from what distance to improve single-cell analysis? In a cell culture, not surprisingly, the involvement of a growing number of neighbours improves accuracy in all of the classifiers until a plateau has been reached (Fig. 3a). As opposed to tissues, we did not observe a tendency, i.e., accuracy did not peak. We assume that this is due to a combined effect of high cellular homogeneity within the cell culture and the fact that in an artificial environment cells do not have the chance to form the characteristic microenvironment during the short plating time. We believe that this tendency can be observed in the majority of cases except where the phenotypes depend on inter-cell communications (such as viral infection spread assays or extracellular vesicle studies).

In the case of the UBC tissue, we found a peak of the optimal distance of neighbours at ∼80-100 micrometres, almost independently of the type of the classification method. The amount of information dramatically increases due to the influence of closer neighbouring superpixels to this point. But those neighbours that are located at further distances, likely due to the presence of other tissue components, led to slowly growing confusion; i.e., the increasing number of neighbouring elements resulted in a decrease in accuracy. In other words, the predictive value of a superpixel decreases with its heterogeneity, which cannot be compensated for by the involvement of more superpixels. A human organ is a complex structure of different tissues. The UBC tissue, which is sufficiently complex in a microscopic field, was selected for our analysis; this field contains at least eight different tissues in a complex but not haphazard manner. The neighbouring superpixels within an optimal distance cover a sufficiently homogeneous area predictive of regular features.

We anticipate that even higher recognition accuracy may be achieved by extending our current work with multi-scale local features that would mimic the pathologist's work using various microscope magnifications to properly understand local and global cellular environment and to draw conclusions regarding any sections examined.

The method presented here can be easily integrated into most single-cell phenotypic analyser pipelines, allowing for the wide-range utilization of its benefit to improve phenotypic characterization at the single-cell level.

## Acknowledgements

P.H. acknowledges support from the Finnish TEKES FiDiPro Fellow Grant 40294/13. L.H. and P.H. acknowledge the European Union and the European Regional Development Funds (GINOP- 2.3.2-15-2016-00001, GINOP-2.3.2-15-2016-00037). The authors thank Dora Bokor PharmD and Gabriella Tick for proofreading the manuscript.

## Author contributions statement

P.H. conceived and led the project. L.H. and F.S. co-supervised the project. T.T. designed the algorithm. N.B., F.K., A.K., T.B., and C.M. developed the software. T.T. prepared the figures. T.T., P.H., L.H., and F.S. designed the experiments and analysed the data. All authors read and approved the final manuscript.

